# RNA-guided endonuclease-driven mutagenesis in tobacco followed by efficient fixation of mutated sequences in doubled haploid plants

**DOI:** 10.1101/042291

**Authors:** Sindy Schedel, Stefanie Pencs, Göetz Hensel, Andrea Müller, Jochen Kumlehn

**Affiliations:** Leibniz Institute of Plant Genetics and Crop Plant Research (IPK), Plant Reproductive Biology, Corrensstr. 3, D-06466 Stadt Seeland/OT Gatersleben, Germany; present address: Martin Luther University, Chair of Plant Breeding, Betty-Heimann-Str. 3, 06120 Halle (Saale), Germany

**Keywords:** genome engineering, haploid technology, pollen embryogenesis, site-directed mutagenesis

## Abstract

**Background:** Customizable endonucleases are providing an effective tool for genome engineering. The resulting primary transgenic individuals are typically heterozygous and/or chimeric with respect to any mutations induced. To generate genetically fixed mutants, they are conventionally allowed to self-pollinate, a procedure which segregates individuals into mutant heterozygotes/homozygotes and wild types. The chances of recovering homozygous mutants among the progeny depends not only on meiotic segregation but also on the frequency of mutated germline cells in the chimeric mother plant.

**Results:** RNA-guided endonuclease-mediated mutagenesis was targeted to the green fluorescent protein gene (*gfp*) harboured by a transgenic tobacco line. Upon retransformation using a *gfp-*specific endonuclease construct, the T_0_ plants were allowed to either self-pollinate, or were propagated via regeneration from *in vitro* cultured embryogenic pollen which give rise to haploid/doubled haploid plants or from leaf explants that form plants vegetatively. Single or multiple mutations were detected in 80% of the T_0_ plants. The majority of these mutations proved heritable by each of the three propagation systems used. Regeneration from *in vitro* cultured embryogenic pollen allowed for homozygous mutants to be produced more efficiently than via sexual reproduction. The recovery of mutations that were not found among sexually produced progeny was shown to be achievable through vegetative plant propagation *in vitro*. In addition, a number of mutations not detected in the primary gRNA/Cas9-expressing plants were uncovered in the progeny, irrespective of the mode of propagation.

**Conclusion:** Regeneration from embryogenic pollen culture provides a convenient method to rapidly generate a variety of genetically fixed mutants following site-directed mutagenesis. Induced mutations that are not sexually transmitted can be recovered through vegetative plant regeneration from somatic tissue.

## Background

Site-specific modifications to genomic DNA sequence, induced by either customizable zinc finger nucleases (ZFNs; Kim et al. 1996), transcription activator-like effector nucleases (TALENs; Christian et al. 2010) or the more recently discovered RNA-guided endonucleases (RGENs), represents the new frontier in genetic engineering. The latter technique relies on a bacterial clustered regularly interspaced short palindromic repeat (CRISPR) loci-encoded defence system (Jinek et al. 2012). All three of the above platforms involve a customizable DNA-binding module (proteinaceous in the case of ZFN and TALEN, and a guide RNA [gRNA] in the case of RGEN) along with a generic enzymatic DNA-cleavage module (*Fok* I for ZFNs and TALENs, *Cas9* for RGENs). The endonuclease’s function is to generate a double strand break (DSB) in, or close to the target site, which is then repaired by either error-prone non-homologous end-joining (NHEJ) or the more precise homology-directed repair (HDR) pathway (Waterworth et al. 2011). Whereas NHEJ tends to induce random insertions, deletions or substitutions at the target DSB site, HDR can be combined with a synthetic repair template to generate predictable sequence modifications.

Effective RGEN-mediated heritable mutagenesis has been demonstrated in both di-and monocotyledonous plant species such as Arabidopsis (Feng et al. 2014, Jiang et al. 2014), rice (Zhang et al. 2014, Zhou et al. 2014), tomato (Brooks et al. 2014), rape seed and barley (Lawrenson et al. 2015). Whereas NHEJ-mediated mutation has been shown in a range of species, so far the use of HDR has been confined to just a few situations (Schiml et al. 2014, Budhagatapalli et al. 2015). The heritability of induced mutations was examined in several plant species with the largely consistent result that not all of the induced mutations are transmitted to the progeny. For example, in a study focussing on *Arabidopsis thaliana*, about 76% of mutations proved to be transmissible (Feng et al. 2014). Primary mutant plants are typically chimeras, so it is necessary to separate and genetically fix mutations by screening the T_1_ generation, a process which is relatively labour-intensive and time-consuming. Selections are then validated in the T_2_ generation. RGEN-mediated mutagenesis in the genus *Nicotiana* was first demonstrated at the cellular level in *Nicotiana benthamiana* (Li et al. 2013, Nekrasov et al. 2013), since when its effectiveness has also been established in *N. tabacum* plants (Gao et al. 2015). However, the inheritance of RGEN-induced mutations in *Nicotiana* spp. has not yet been demonstrated.

The present study details a number of heritable RGEN-induced mutations to the *gfp* transgene used here as experimental target in tobacco. It aims to demonstrate that induced mutations are heritable and that genetic fixation of altered sequences can be particularly efficiently achieved by regeneration of progeny plants from *in vitro* cultured embryogenic pollen. In addition, an attempt was made to use vegetative propagation for the recovery of mutations which are not found in progeny derived from self-fertilization.

## Results

### gRNA/Cas9-mediated genetic modification of gfp

A T-DNA construct comprising a *gfp-*specific gRNA, *Cas9* and *bar* (encoding phosphinothricin acetyltransferase, the activity of which ensures resistance to the herbicide Bialaphos) was introduced into a transgenic tobacco line harbouring a single copy of *gfp*, along with *NptII* (encoding neomycin phosphotransferase). When the presence of the *Cas9* and the *gRNA* sequences in the genomic DNA extracted from Bialaphos-resistant regenerants was tested by PCR (primer pairs detailed in Additional File 1), 15 of the 21 regenerants were shown to harbour both transgene sequences. The *gfp* region in these 15 individuals was then characterized using the T7 endonuclease (T7E1) assay. The 561 bp *gfp* amplicons were melted and re-annealed, in order to produce a heteroduplex where more than one variant of the sequence was present; this structure is cleavable by T7E1. The T7E1 profile of the amplicons derived from 12 of the 15 gRNA/Cas9-positive plants comprised two fragments (274 and 287 bp), which resolved as a single band following 1.5% agarose gel electrophoresis (Fig. 1a). By contrast, none of the six transgenic plants lacking Cas9 and/or gRNA scored positive with respect to the T7E1 assay. The sequencing of individual clones derived from the *gfp* amplicons of five of the 12 mutation-carrying plants (#122, #125, #254, #258 and #264) revealed the presence of deletions (ranging in length from 1-91 bp) or insertions of 1 bp (Fig. 1b). Across all T_0_ plants examined, 62.5% of the independently generated mutations caused a translational reading frame shift, and 60% of these frame shifts entailed the formation of a premature stop codon. Some of the deletions (corresponding to a loss of 2-18 amino acids in the gene product) did not induce a reading frame shift. Plants #122 and #125 both harboured more than one mutated *gfp* sequence, while the other three plants only harboured a single one. In all five plants, a wild type (WT) *gfp* sequence was retained, indicating that the plants were either heterozygous and/or chimeric.

**Fig. 1.**
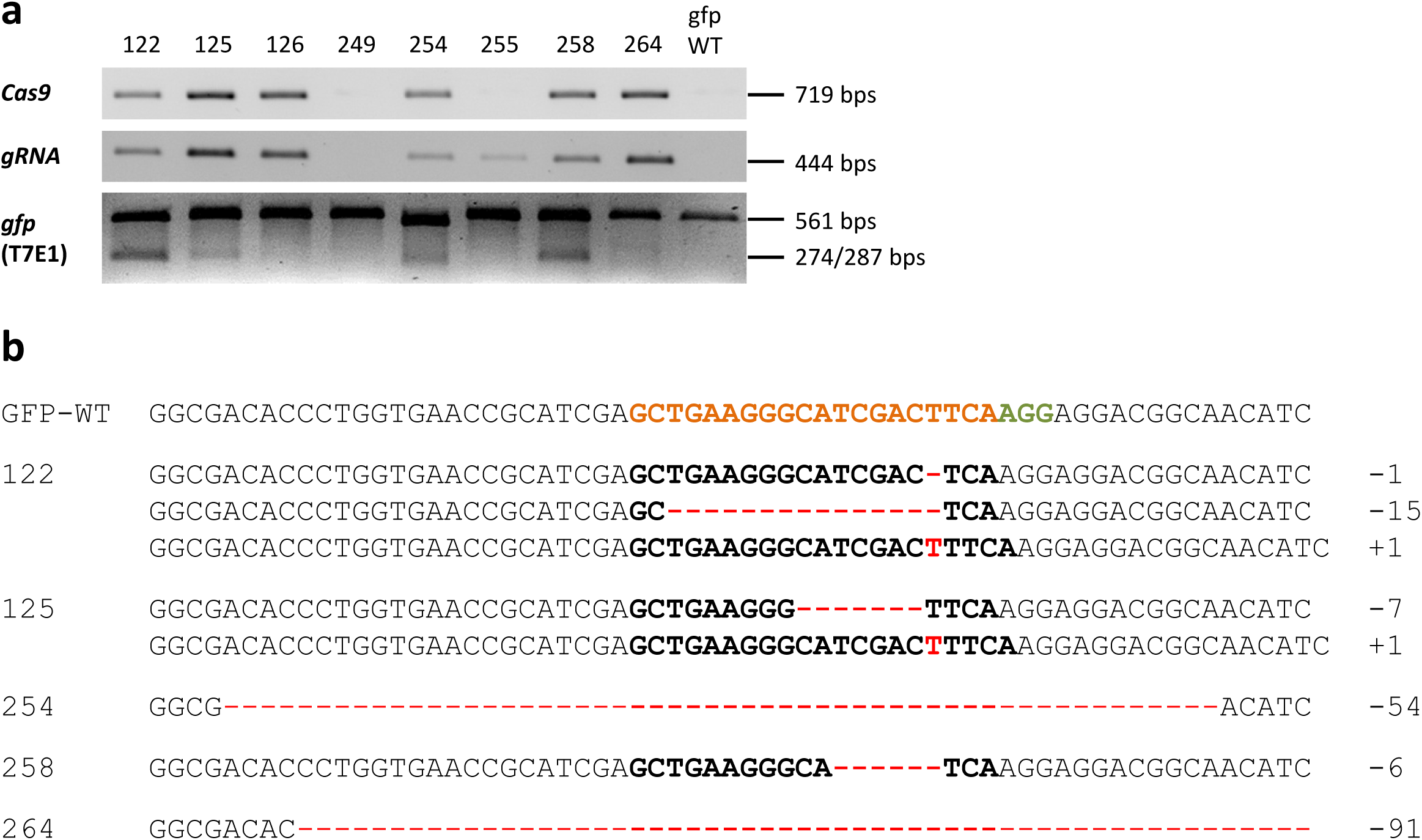
Modifications of the *gfp* sequence induced by RGEN. (a) The presence of *Cas9* and *gRNA* and induced mutations to *gfp* in the T_0_ generation. The first two rows show the outcomes of PCRs targeting *Cas9* and *gRNA* and the wild type *gfp* sequence (gfp-WT). The lower row shows the outcome of the T7E1 assay. The expected cleavage products were of length 267 and 294 bp, which resolve electrophoretically as a single band. (b) Sequencing outcomes of individually cloned *gfp* sequences from five T_0_ plants. The numbers of induced nucleotide changes are indicated to the right of each sequence. The sequence coloured orange is the protospacer sequence and the one coloured green is the protospacer-adjacent motif; deletions are represented by dashes, insertions by red letters.

**Additional file 1.**
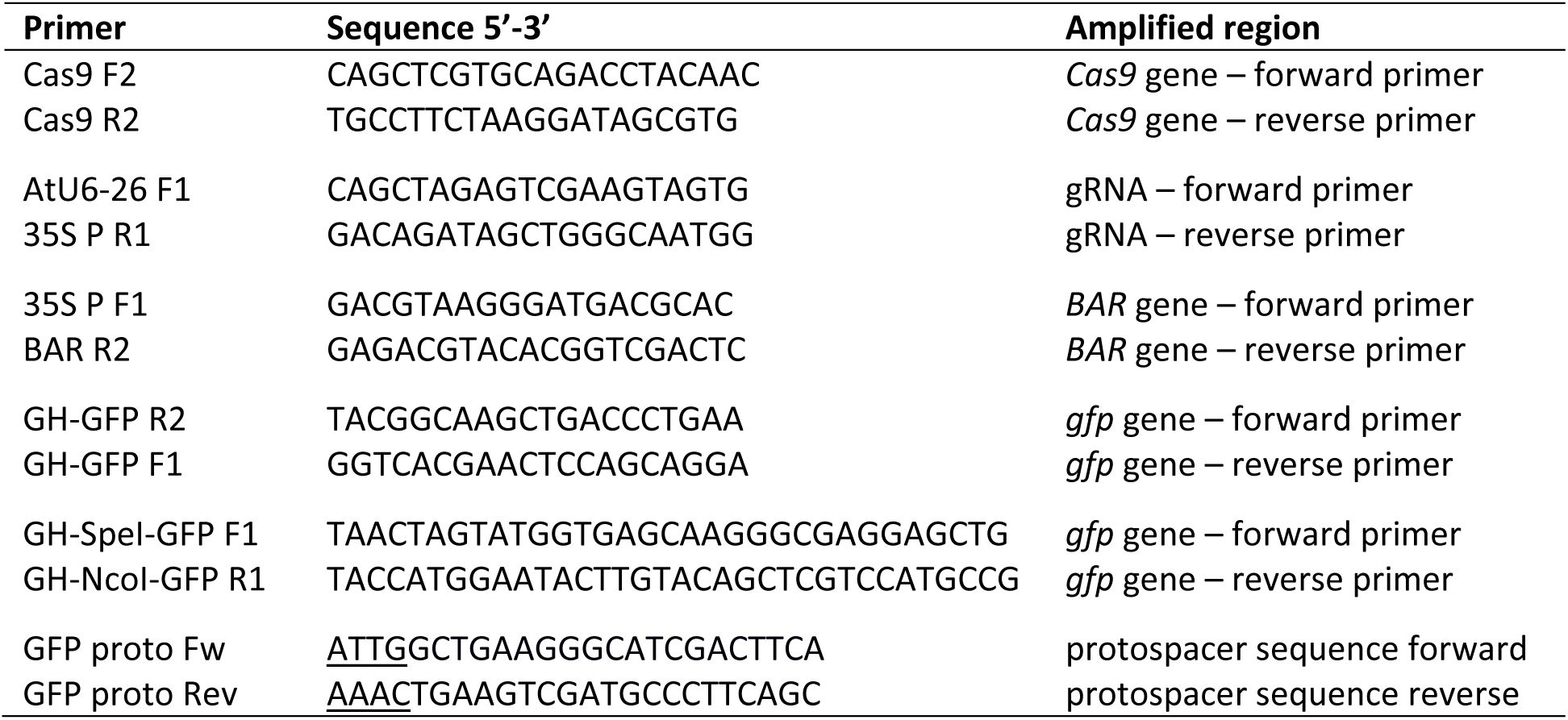
Primers used for the identification of T-DNA regions, the amplification of the *gfp* target region and the cloning of binary vectors pGH292 and pSI24. Oligonucleotides used for the identification of T-DNA regions, the amplification of the *gfp* target region and the cloning of binary vectors pGH292 and pSI24.

### Sexual transmission of the gfp mutations

T_1_ progeny of plants #125 and #254 (each carrying multiple *gfp* mutations), and #126 (harboring no detectable *gfp* mutation despite the presence of *gRNA* and Cas9) was analysed in detail. Between 15 and 50 progeny per selfed T_0_ plant were subjected to the T7E1 assay, and the PCR products of those samples flagged as negative were sequenced directly in order to discriminate between WT and mutant homozygotes. Several of the T7E1-positive progeny (i.e., those which harboured two or more *gfp* variants) were examined in more detail by generating individual clones from their *gfp* amplicon, and then sequencing these. The T_1_progenies were shown to include some non-WT individuals in each case (Fig. 2, Table 1). Among the 20 T_1_ offspring of plant #125 analyzed in detail, the 1 bp insertion segregated, but the 7 bp deletion was not recovered among the progeny generated via selfing. Four deletions (1, 3, 6 and 24 bp) were uncovered in the T_1_ generation even though they had not been detected in the mother plant. In plant #254, the 54 bp deletion proved to be heritable, and a 132 bp deletion was additionally revealed in the T_1_ generation. In all, two thirds of the *gfp* mutations detected in T_0_ plants #125 and #254 were transmissible, while 71.4% of all independently generated mutations detected across both generations were first found in T_1_plants. Mutations were also uncovered among the progeny of plant #126: specifically, three micro-deletions (1, 3 and 4 bp) and the identical 1 bp insertion that was independently induced in the T_0_ plant #125.

**Fig. 2.**
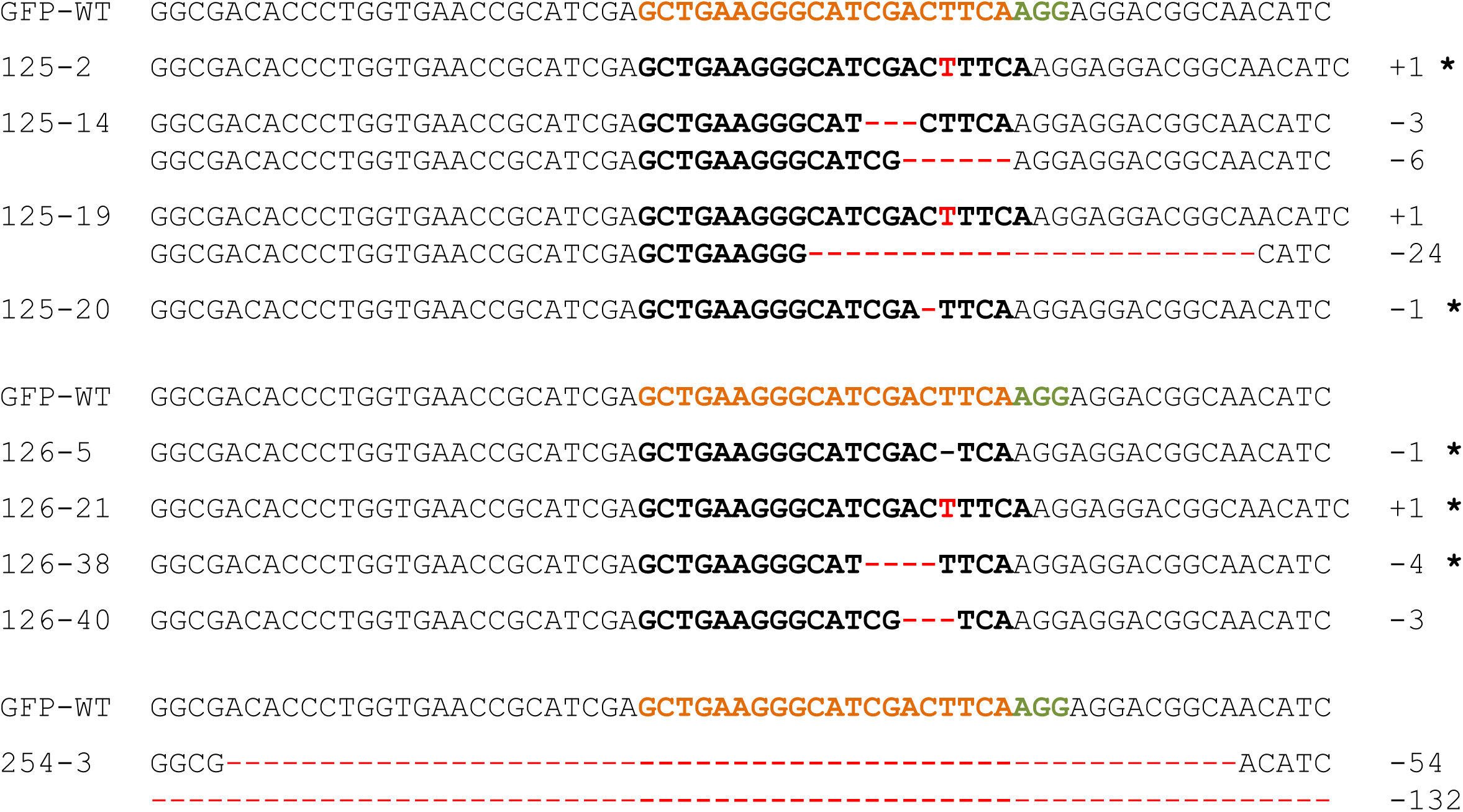
The inheritance of targeted mutagenesis products. Alignment of the *gfp* sequences of descendants of T_0_ plants #125, #126 and #254. The numbers of nucleotides deleted and inserted are shown to the right of each sequence. The sequence coloured orange is the protospacer sequence and the one coloured green is the protospacer-adjacent motif; deletions are represented by dashes, insertions by red letters.

**Table 1.** Analysis of progeny produced via selfing and haploid technology. The frequency of mutations among progeny of the T_0_ mother plants produced by either self-fertilization or passage through embryogenic pollen culture.

|  | Selfing |  |  |  | Haploid technology |
| --- | --- | --- | --- | --- | --- |
| T<sub>0</sub> plant identifier | 125 | 126 | 254 | average | 125 |
| no. of mutations found in T<sub>0</sub> | 2 | 0 | 1 | 1.0 | 2 |
| no. of T<sub>1</sub> plants analyzed | 50 | 50 | 15 | 38.3 | 62 |
| non-mutant T<sub>1</sub> plants | 5 (10%) | 11 (22%) | 4 (26.7%) | 6.7 (17.5%) | 0 |
| T<sub>1</sub> mutant plants | 45 (90%) | 39 (78%) | 11 (73.3%) | 31.6 (82.5%) | 62 (100%*) |
| heterozygous/chimeric T<sub>1</sub> mutant plants | 28 (56%) | 29 (58%) | 11 (73.3%) | 22.6 (59%) | 25 (40.3%) |
| genetically fixed T<sub>1</sub> mutant plants | 17 (34%) | 10 (20%) | 0 | 9.0 (23.5%) | 37 (59.7%) |
| no. of different mutations found in T<sub>1</sub> | 5 | 4 | 2 | 3.6 | 12 |
| a) no. of mutations found in T<sub>0</sub> and T<sub>1</sub> | 1 | - | 1 | 0.6 | 2 |
| not genetically fixed in T<sub>1</sub> plants | 0 | - | 1 | 0.3 | 0 |
| genetically fixed in T<sub>1</sub> plants | 1 | - | 0 | 0.3 | 2 |
| b) no. of mutations not found in T<sub>0</sub> but found in T<sub>1</sub> | 4 | 4 | 1 | 3.0 | 10 |
| not genetically fixed in T<sub>1</sub> plants | 3 | 1 | 1 | 1.7 | 3 |
| genetically fixed T<sub>1</sub> plants | 1 | 3 | 0 | 1.3 | 7 |

\*) pollen embryogenesis was conducted under Bialaphos-based selection for RGEN-transgenics

*) pollen embryogenesis was conducted under Bialaphos-based selection for RGEN-transgenic

In all, 54% of #125 T_1_ progeny carried one or more mutated *gfp* sequence in addition to the WT *gfp* allele, 36% were homozygous for a mutant *gfp* allele and the remaining 10% inherited no *gfp* mutation. Of the plant #254 T_1_ progeny analysed, 73.4% were heterozygous/chimeric plants and the other 26.6% carried only the WT *gfp* allele. Finally, 58% out of the plant #126 T_1_ progeny were heterozygous and/or chimeric, 20% were homozygous for a mutant *gfp* allele and the remaining 22% inherited no *gfp* mutation. Among the genetically fixed progeny, four lacked the RGEN-coding T-DNA (as shown by a PCR directed at *Cas9*) but had retained the identical 1 bp insertion. Of the heterozygous/chimeric T_1_ progeny, 21 also lacked *Cas9*.

### Efficient production of non-chimeric, homozygous mutants by means of pollen embryogenesis

In an attempt to accelerate the recovery of non-chimeric, homozygous mutants from chimeric #125 T_0_ mother plant, an embryogenic pollen culture was initiated. All of the 62 Bialaphos-resistant regenerants produced via pollen embryogenesis retained the RGEN T-DNA. An analysis based on the T7E1 assay and the direct sequencing of T7E1-negative PCR products revealed that every plant carried at least one altered *gfp* sequence (Table 1). In 59.7% of all examined plants produced via pollen embryogenesis, only single mutated *gfp* sequences were detected (associated with negative outcome in the T7E1 assay) and the WT *gfp* sequence was absent, suggesting the successful fixation of the mutant sequence. Both of the mutations present in the T_0_ generation (a 1 bp insertion and a 7 bp deletion) were represented among these 37 regenerants, along with ten additional mutations: these included a 2 bp substitution (plant #125-DH44; Fig. 3). The remaining 40.3% of the regenerants were positive for the T7E1 assay, indicating their retention of at least two *gfp* sequences. The sequencing of individual clones of the *gfp* amplicon produced from five of these 25 regenerants revealed that in four, a single altered *gfp* sequence was accompanied by the WT sequence, while in plant #125-DH13 there were three distinct non-WT sequences in addition to the WT one (Fig. 3); one of the variants involved both a 6 bp deletion and a 1 bp insertion.

**Fig. 3.**
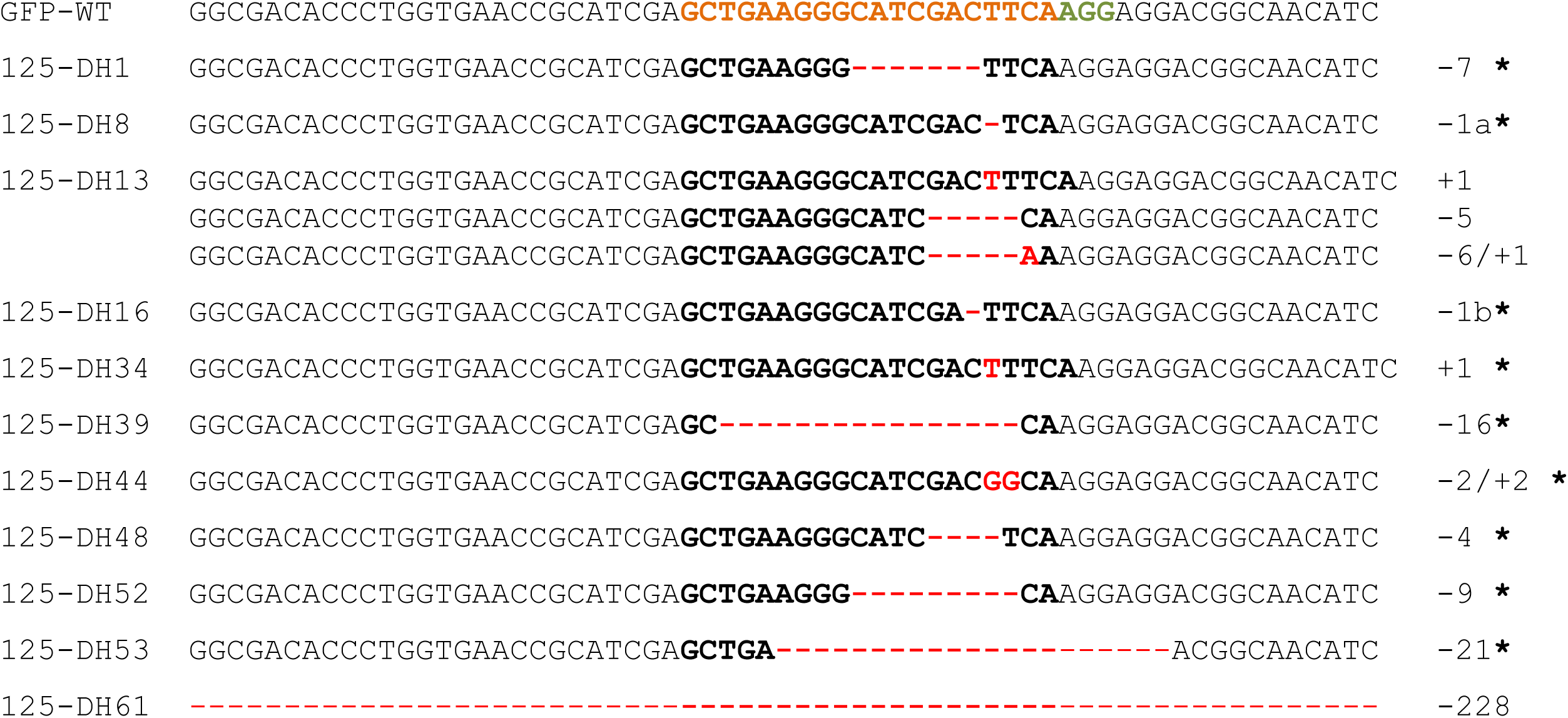
Separating mutated sequences by regeneration from embryogenic pollen culture. The sequences of various regenerants derived from plant #125 are shown. The numbers of nucleotide changes are indicated to the right of each sequence. The sequence coloured orange is the protospacer sequence and the one coloured green is the protospacer-adjacent motif; deletions are represented by dashes, insertions by red letters.

### Maintaining mutations via in vitro vegetative propagation

Both the timing and location of mutagenesis affect whether or not a mutation will be transmitted to the next generation. Not all of the mutations detected in the T_0_ plants were transmitted to their progeny, as shown above. Thus, an attempt was made to recover some of these mutations via vegetative propagation. Leaf segments from the T_0_ plant #125 were cultured *in vitro*, from which 15 Bialaphos-resistant regenerants were recovered. A T7E1 assay of their *gfp* amplicon indicated the retention of more than one *gfp* variant in 13 of the 15 plants. The sequencing of individual clones from the *gfp* amplicon produced from four of these showed that in one case (plant #125-SC5) there remained only a single altered sequence (the 7 bp deletion) along with the WT *gfp* sequence, while in the other three, multiple variants were accompanied by the WT sequence. The latter mutations included the maternal 1 bp insertion, along with some additional ones, notably the 92 bp deletion plus a 2 bp insertion present in plant #125-SC1 (Fig. 4). The direct sequencing of the PCR products derived from the two regenerants which were negative for the T7E1 assay revealed the absence of the WT *gfp* sequence along with a single *gfp* variant in both cases, indicating the likely fixation of the mutant sequence. Overall, 86.7% of the regenerants analysed were heterozygous or chimeric for a mutation, while the remainder were homozygous. Importantly, both the 1 bp insertion and the 7 bp deletion detected in the T_0_ plant were maintained, and seven additional mutations occurred (Fig. 4).

**Fig. 4.**
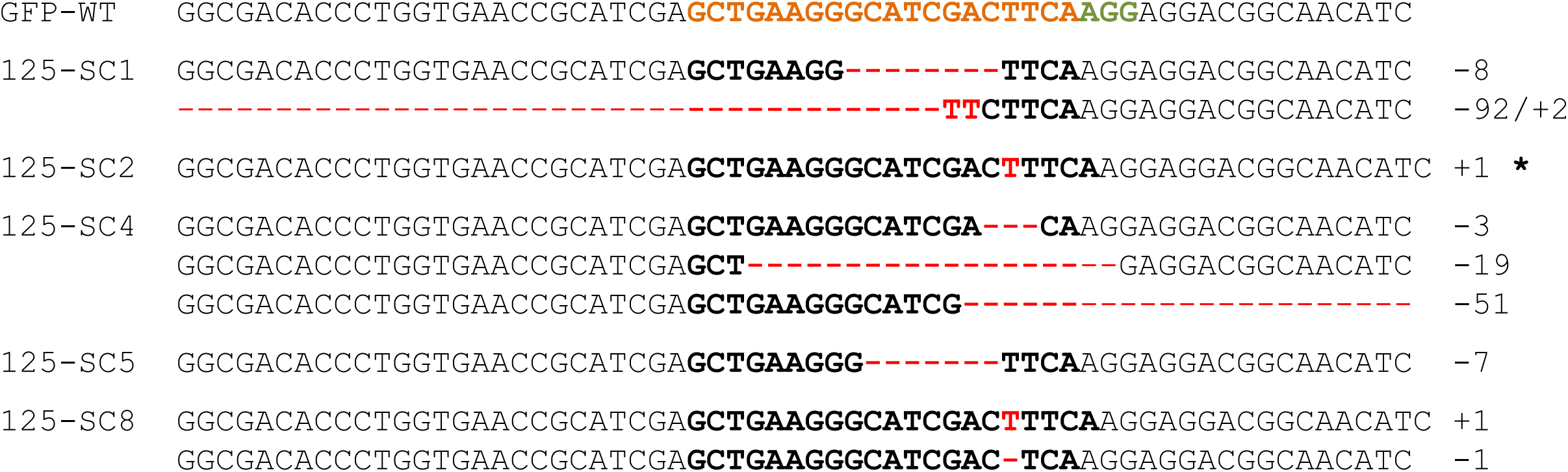
RGEN-induced mutations in the *gfp* sequence among regenerants derived from leaf explants from T_0_ plant #125. The figure shows an alignment of the *gfp* sequences recovered. The number of nucleotides deleted (dashes) and inserted (red letters) are shown to the right of each sequence. The sequence coloured orange is the protospacer sequence and the one coloured green is the protospacer-adjacent motif.

### Variety and frequency of mutations

To investigate the variety of induced mutations and the frequency of independent formation of specific alterations, the data derived from each of the four sets of plant material (T_0_ plants, conventional T_1_ progeny, and regenerants from both embryogenic pollen culture and leaf explants) were combined. A total of 32 distinct mutation events to the *gfp* sequence was revealed; of these 26 (81.2%) were deletions, three (9.4%) were insertions, one (3.1%) was a 2 bp substitution and two (6.2%) involved an insertion of 1 or 2 bp in conjunction with a deletion. Half of the deletions were shorter than 10 bp, and the largest was 228 bp long; all of the insertions not accompanied by a deletion were of 1 bp in length. Overall, 25% of the mutations involved just a single base pair (Fig. 5). No reading frame shift was induced in 43.8% of the mutations: these included some deletions longer than 2 bp, the 2 bp substitution in plant #125-DH44 and the combined 92 bp deletion / 2 bp insertion event in plant #125-SC1.

**Fig. 5.**
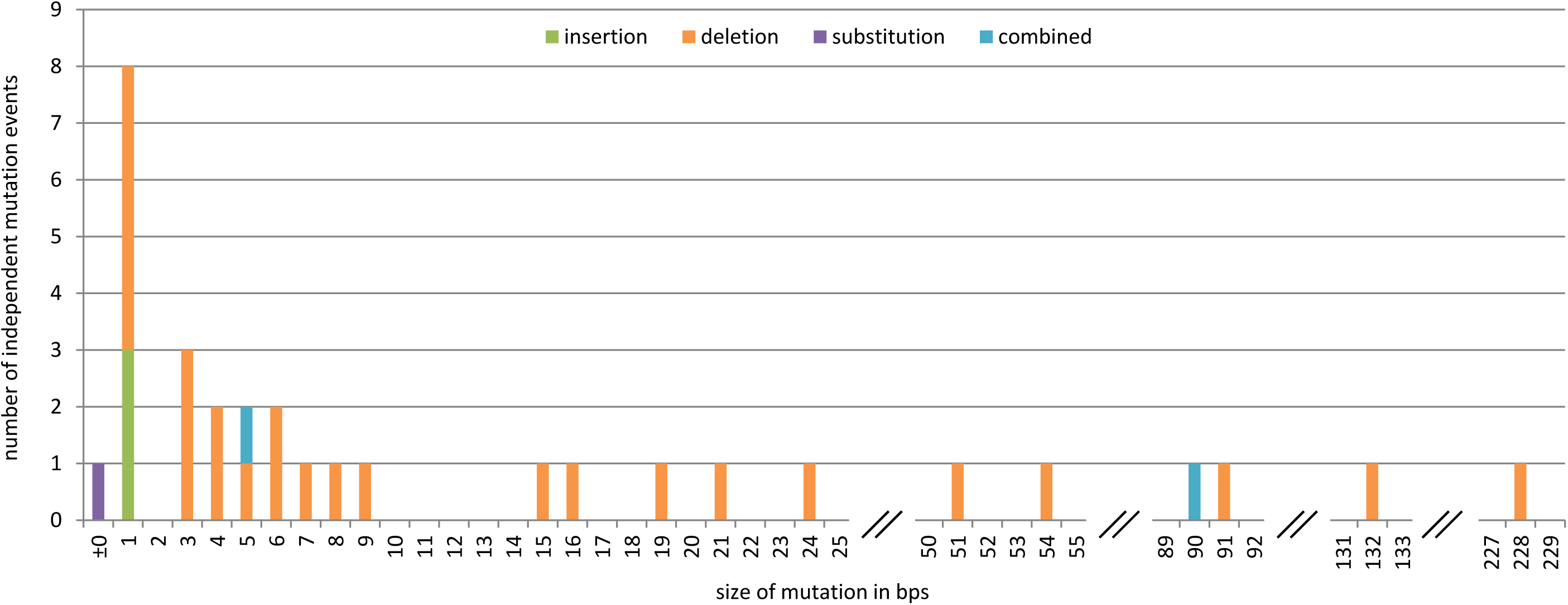
The variety of induced *gfp* mutations and the frequency of types and lengths of alterations. The *x* axis plots the extent of the sequence alteration in bp and the *y* axis their frequency. Identical mutations were considered when they derived from independent events.

## Discussion

Introducing a *gfp-*specific *gRNA/Cas9* construct into tobacco resulted in a high proportion (80%) of the T_0_ plants (carrying both gRNA and Cas9 expression units) experiencing one or more mutations to the target locus. A similar level of success has been reported in the same species, although targeting two different genes (*PDS* and *PDR6*) (Gao et al. 2014).

The considerable frequency with which the T-DNA was only partially integrated (in 28.6% of the T_0_ plants) in the present study may reflect the instability of its left border sequence, since the right border sequence is known to be integrated more precisely than the left one (Tinland 1996, Kim et al. 2003). Whereas placing the selectable marker gene adjacent to the left T-DNA border is thought to result in a high proportion of transgenics carrying the complete T-DNA, the vector used in the present study carried the selectable marker at the right border end of the T-DNA. In the few documented cases where the integrity of the *Cas9* and *gRNA* containing T-DNA has been investigated, some partial integrations were also observed, and, expectedly, the respective plants proved non-mutant (Zhou et al. 2014, Xu et al. 2015).

Sexual transmission of targeted mutations has not been demonstrated in a *Nicotiana* sp. before. In the present study, the transmission of the mutations to the T_1_ generation mirrors that experienced in both rice (Zhou et al. 2014, Xu et al. 2015) and *A. thaliana* (Feng et al. 2014, Jiang et al. 2014), in that not every mutation revealed in the T_0_ plant was recoverable. For example, the 1 bp insertion present in T_0_ plant #125 was represented in the sample of 50 T_1_ progeny examined, but the 7 bp deletion was not (Fig. 2). However, the latter event was maintained both via pollen embryogenesis and *in vitro* vegetative propagation (Fig. 4). Consequently, both of these alternative ways of producing progeny can be considered useful for the recovery of mutations that are too rarely or not at all transmitted sexually.

In addition, mutations that had not been detected in #125 were uncovered among progeny of this plant, irrespective of the mode of propagation. However, it is not known which of these mutations had been present in the T_0_ plant but remained undetected and which ones were newly induced in progeny plants which had inherited the gRNA and Cas9 expression units along with the *gfp* WT allele. Notably, T_0_ plant #126 appeared lack any mutations, yet some of its T_1_ progeny harboured mutants, even in the homozygous state (Fig. 2). The homozygosity of almost half of these new mutants in the T_1_ generation (Table 1) suggests the mutations were likely to have been induced in the T_0_ plant without being uncovered for the time being. Note that Zhang et al. (2014) have detected various mutations when DNA prepared from different parts of a T_0_ plant was compared. Two possible scenarios present themselves: first, that the mutations were induced either after the DNA had been sampled or occurred elsewhere in the T_0_ plant; and/or secondly, that too small proportion of the T_0_ plant’s DNA analysed harboured the altered *gfp* sequence(s) for the T7E1 assay to pick the effect up. The fixation of mutations independent of passage through meiosis cannot, however, be ruled out, given that two of the regenerants from vegetative propagation carried a mutation in the homozygous condition. A possible explanation for this rather unexpected outcome is that a DSB was induced by the gRNA-guided *Cas9* in the WT *gfp* allele still present in a heterozygous mutant cell, and then was repaired via homologous recombination, recruiting the previously altered sequence as repair template.

A relatively high percentage - perhaps even all - of the T_0_ plants were chimeric with respect to mutations at the target site, which is possible not only given that a regenerating shoot can originate from more than one cell (Schmuelling & Schell 1993, Li et al. 2009), but also because RGENs in principle can generate a number of independent mutation events in various cells of a developing individual as long as an intact target allele is present. The resolution of chimeras, as in any mutagenesis programme, normally requires passage through meiosis, which can be achieved either via conventional gamete fusion, or, more efficiently, by exploiting the totipotency of immature pollen. While ~60% fixation was achieved by regenerating plants from embryogenic pollen culture, the rate was only half of this among the conventionally generated T_1_ progeny (Table 1). Note that no homozygous mutants were obtained by the conventional route from plant #254. Similar difficulties have been experienced in fixing mutations induced in *A. thaliana* (Feng et al. 2014).

The variety and frequency of induced mutations described here are consistent with the experience of RGEN applied to rice, in which a high frequency of small deletions can be induced, along with 1 bp insertions only (Zhang et al. 2014). In *A. thaliana*, it is however, rather different, since a range in insertion lengths is the outcome (Feng et al. 2014). The basis of this apparent contradiction may lie in species-specific differences in the activity and preferences of the complex endogenous DNA repair mechanisms.

## Conclusion

The majority of site-directed mutations induced by RGENs in tobacco proved to be transmissible, irrespective of whether propagation was via self-fertilization or the *in vitro* culture of either embryogenic pollen or somatic tissue. Regeneration from embryogenic pollen culture proved particularly effective as a tool to rapidly generate genetically fixed mutant plants, while at least some of the mutations, which appear to not be transmitted through the germline, can be maintained via vegetative propagation.

## Material and Methods

### Plant growth

Seeds of the wild type *Nicotiana tabacum* accession SR1 and the single *gfp* transgene copy line TSP20 1–1 were surface-sterilized and germinated for two weeks on solidified Murashige and Skoog (MS) medium (Murashige & Skoog, 1962). They were subsequently transferred into boxes (107 x 94 x 96 cm) containing MS medium and left to grow for 6-8 weeks. Once regenerants from tissue culture had developed a viable root system, they were potted into soil and grown under a 16 h photoperiod provided by 35,000 lux light at 22/20°C, then re-potted and grown for a further ~10 weeks (20/18°C, 16 h photoperiod, 30,000 lux) to obtain progeny by self-fertilization. T_1_ seed was germinated in soil under 22/20°C day/night, 16 h photoperiod (35,000 lux) light.

### T-DNA constructs

The *gfp* sequence was amplified using the primer pair GH-SpeI-GFP F1/GH-NcoI-GFP R1 (Additional file 1) and introduced into the pCR2.1 plasmid (Invitrogen, Carlsbad, CA, USA) to form pGH124. A *gfp-*containing *Spe*I/*Eco*RI fragment of pGH124 was inserted into pNos-AB-M (DNA Cloning Service, Hamburg, Germany) to yield pGH119. Subsequently, the *Spe*I/*Hin*dIII *gfp*-containing fragment was subcloned into pUbiAT-OCS (DNA Cloning Service) between the *A. thaliana UBIQUITIN-10* promoter and the *Agrobacterium tumefaciens OCS* termination sequence, the resultant plasmid being denoted pGH167. An *Sfi*I fragment of pGH167 harbouring the entire *gfp* expression cassette was integrated into the binary vector pLH9000 (DNA-Cloning-Service) to produce pGH292, which was introduced into *A. tumefaciens* strain GV2260 via a heat shock protocol.

The Gateway^®^-compatible RGEN expression system (Fauser et al. 2014) was used to construct a *gfp*-specific derivative. In the first step, the *gfp*-specific protospacer sequence (annealed oligonucleotides, Additional file 1) was introduced into pEN-Chimera by exploiting its two *Bbs*I sites. The resulting gRNA-encoding chimera, driven by the *A. thaliana U6-26* promoter, was then transferred into pDe-CAS9 through a single site Gateway^®^ LR reaction to form an expression cassette where the *Cas9* (codon optimized for *A. thaliana*) was driven by the *Petroselinum crispum Ubi 4-2* promoter and its terminator sequence was *pea3A* from *Pisum sativum*. The resulting binary vector (pSI24) was introduced into Agrobacterium as described above.

### Tobacco transformation

Fully developed leaves of 6-8 week old plants were cut into ~1 cm^2^ pieces and cultured on MS medium containing 3% (w/v) sucrose, 1 mg/L 6-benzylaminopurine, 0.1 mg/L 1- naphthalene acetic acid and 2% (w/v) bacto agar. After one or two days, the leaf segments were bathed in a suspension of Agrobacterium cells (OD_600_ 0.2) for 30 min, blotted dry and then replaced on the culture medium and held in the dark at 19°C for three days. The leaf segments were subsequently removed to the same medium containing 400 mg/L Ticarcillin and either 100 mg/L kanamycin in the case of transformation with GV2260/pGH292 or 5 mg/L Bialaphos when GV2260/pSI24 was used, and sub-cultured every ten days (initially in the dark, then in the light) at 22°C. Differentiated shoots were separated from the callus and transplanted into a root induction MS-based medium containing 2% (w/v) sucrose, 0.8% (w/v) bacto agar and antibiotics (as described above). Rooted plantlets were later transferred to soil and grown in a greenhouse to maturity.

### Vegetative in vitro propagation from leaf explants

In order to vegetatively propagate the T_0_ plant #125, explants from different leaves were surface-sterilized by immersion first in 70% ethanol and then in 5% (w/v) sodium hypochlorite, then rinsed three times in sterile water. A number of ~1 cm^2^ pieces was cut and placed on MS medium containing 3% (w/v) sucrose, 1 mg/L 6-benzylaminopurine, 0.1 mg/L 1-naphthalene acetic acid, 2% (w/v) bacto agar, 5 mg/L Bialaphos and 400 mg/L Ticarcillin. Regeneration was handled as for the transgenic material, as described above.

### Embryogenic pollen culture

The procedure used for embryogenic pollen culture followed Floss et al. (2009). The selection of regenerants was based on the presence in the medium of 5 mg/L Bialaphos.

### Genotyping

Genomic DNA was isolated from leaf tissue using a phenol chloroform-based procedure (Palotta et al. 2000) to serve as a template for PCRs targeting *bar*, *Cas9* and *gRNA*, employing primer pairs as listed in Additional file 1. To detect mutations in *gfp*, the genomic region surrounding the target sequence was PCR amplified using the primer pair GH-GFP R2/F1 (Additional file 1), and the amplicons, following their purification using a QIAquick PCR purification kit (QIAGEN, Hilden, Germany), were subjected to the T7E1 assay (NEB, Ipswich, MA, USA). PCR products from T_1_ plants scored as negative were sequenced directly, while those which were T7E1-positive were cloned into the pGEM^®^-T Easy vector (Promega, Madison, WI, USA) following the manufacturer’s protocol and resultant clones were individually sequenced.

## Acknowledgement

We thank H. Puchta (KIT Karlsruhe) for the gift of the Gateway^®^-compatible *Cas9* expression system.

